# A specialist flea beetle manipulates and tolerates the activated chemical defense in its host plant

**DOI:** 10.1101/2021.03.12.435088

**Authors:** Theresa Sporer, Johannes Körnig, Natalie Wielsch, Steffi Gebauer-Jung, Michael Reichelt, Yvonne Hupfer, Franziska Beran

## Abstract

Glucosinolates, the characteristic secondary metabolites of Brassicales, are hydrolyzed upon herbivory by myrosinases to toxic and deterrent defense metabolites. The specialist flea beetle, *Phyllotreta armoraciae*, sequesters glucosinolates in the body despite myrosinase activity, but it is unknown whether plant myrosinase activity influences sequestration and how beetles prevent the hydrolysis of ingested glucosinolates. In feeding experiments performed with the myrosinase-deficient *Arabidopsis thaliana tgg1×tgg2* (*tgg*) mutant and the corresponding wild type, we found that plant myrosinases reduced the glucosinolate sequestration rate by up to 50% and hydrolyzed a fraction of ingested glucosinolates in adult beetles. Although these results show that *P. armoraciae* cannot fully prevent glucosinolate hydrolysis, we observed no negative influence on beetle performance. To understand how *P. armoraciae* can avoid the hydrolysis of some ingested glucosinolates, we analyzed their fate directly after ingestion. *P. armoraciae* rapidly absorbed glucosinolates across the gut epithelium, a strategy that has been proposed to prevent hydrolysis in the gut lumen of sequestering insects. Moreover, beetle gut content suppressed *in vitro* myrosinase activity, and almost no myrosinase activity was detectable in the feces, which indicates that ingested myrosinases are inactivated in the beetle gut. In summary, we show that *P. armoraciae* uses several strategies to prevent the hydrolysis of ingested glucosinolates but can also tolerate the formation of glucosinolate hydrolysis products.

## Introduction

Many plants deter herbivores with a chemical defense that is activated upon tissue damage (Morant et al., 2008). The chemical defense compound is usually stored as inactive glucose conjugate in the vacuole of plant cells. When plant tissue is damaged, the glucose moiety is hydrolyzed by a defensive β-glucosidase, originally localized separately from the glucose conjugate, which liberates toxic and deterrent compounds (Morant et al., 2008; Pentzold et al., 2014b). Activated chemical defenses are usually more efficient in deterring chewing herbivores that cause extensive tissue damage than in deterring insects with less invasive feeding modes, such as sap suckers (Pentzold et al., 2014b).

A number of herbivorous insects evolved resistance against this plant defense strategy (Pentzold et al., 2014b) and some highly adapted species even accumulate (sequester) plant glucosides in their bodies and deploy them for defense against predators (Kazana et al., 2007; Opitz and Müller, 2009; Beran et al., 2019). However, it is currently not well understood how chewing insects prevent the hydrolysis of ingested plant glucosides by plant β-glucosidases. A study with turnip sawfly larvae suggests that a rapid absorption of ingested glucosides (glucosinolates) across the gut epithelium prevents their hydrolysis in the gut; this is possibly facilitated by low plant β-glucosidase activity in the anterior gut (Abdalsamee et al., 2014). A rapid uptake mechanism was also proposed to enable western corn rootworm larvae, *Diabrotica virgifera virgifera*, to sequester benzoxazinoid glucosides; however, there was no evidence for reduced β-glucosidase activity in the larval gut (Robert et al., 2017). Burnet moth larvae which sequester cyanogenic glucosides avoid extensive plant β-glucosidase activity by a leaf-snipping feeding mode causing only minor tissue damage. Moreover, the common alkaline pH in the midgut lumen of lepidopteran larvae also inhibits plant β-glucosidase activity, whereas saliva extracts of burnet moth larvae did not prevent cyanogenic glucoside breakdown (Pentzold et al., 2014a).

Sequestering herbivores must avoid the hydrolysis of the ingested plant glucosides they sequester, but the extent to which plant β-glucosidase activity influences glucoside sequestration has rarely been assessed. The brassicaceous model plant *Arabidopsis thaliana* offers an ideal system to address this question. The activated defense of *Arabidopsis* and other plants of the order Brassicales is the glucosinolate-myrosinase system (Halkier and Gershenzon, 2006; Blažević et al., 2020). Glucosinolates are a structurally diverse group of amino acid-derived thioglucosides that are hydrolyzed by β-thioglucosidases called myrosinases. The resulting aglucone is unstable and rearranges spontaneously into a highly reactive isothiocyanate, which is toxic for small herbivores (Jeschke et al., 2016a). The *Arabidopsis tgg1×tgg2* double knock-out mutant mutant (*tgg*) is devoid of myrosinase activity in leaves (Barth and Jander, 2006) and thus can be used for comparative feeding studies with the corresponding *Arabidopsis* Col-0 wild type (wild type). For example, in feeding experiments performed with the cabbage stem flea beetle, *Psylliodes chrysocephala*, adult beetles sequestered six times more glucosinolates from the myrosinase-deficient *Arabidopsis tgg* mutant than from the wild type (Beran et al., 2018). In a similar feeding experiment performed with larvae of the horseradish flea beetle, *Phyllotreta armoraciae*, only traces of sequestered glucosinolates were detected in wild type-fed larvae, whereas comparatively high glucosinolate levels were found in *tgg*-fed larvae (Sporer et al., 2020). In contrast to *P. armoraciae* larvae, adult beetles were able to sequester glucosinolates from wild type leaves (Yang et al., 2020), which suggests that plant myrosinase activity has a stronger influence on glucosinolate sequestration in larvae compared to adults. Plant myrosinase activity influences not only sequestration, but also the feeding behavior of *Phyllotreta* flea beetles. In field experiments, the crucifer flea beetle, *Phyllotreta cruciferae*, caused less feeding damage on *Brassica rapa* plants selected for high myrosinase activity than on plants selected for low myrosinase activity (Siemens and Mitchell-Olds, 1996).

Here, we investigated the influence of plant myrosinase activity on glucosinolate sequestration in the adult life stage of *P. armoraciae*. In a previous feeding study, we recovered about 35% of the total ingested glucosinolates from wild type leaves in the beetle body and feces, whereas the metabolic fate of more than 60% of the total ingested glucosinolates remained unknown (Yang et al., 2020). One possible explanation is that the unrecovered glucosinolates were hydrolyzed by plant myrosinases. To investigate this possibility, we performed a series of comparative feeding experiments with myrosinase-deficient and wild type *Arabidopsis* plants. In nature, *P. armoraciae* is closely associated with horseradish, *Armoracia rusticana*, a plant species that is characterized by high levels of allyl glucosinolate (Li and Kushad, 2004; Ciska et al., 2017). Therefore, we additionally investigated the influence of plant myrosinase activity on the sequestration of allyl glucosinolate by spiking the intact glucosinolate into *Arabidopsis* leaves. We observed a negative influence of plant myrosinase activity on glucosinolate sequestration and confirmed that a fraction of ingested glucosinolates is hydrolysed in *P. armoraciae*. We thus asked whether the metabolism of glucosinolate hydrolysis products incurs a metabolic cost in *P. armoraciae*. Finally, we explored possible mechanisms that allow *P. armoraciae* to partially prevent the hydrolysis of ingested glucosinolates by plant myrosinase.

## Materials and Methods

### Plants and insects

Food plants, *Brassica juncea* cv. “Bau Sin” and *Brassica rapa* cv. “Yu-Tsai-Sum” (Known-You Seed Co., Ltd., Taiwan) were cultivated in a controlled environment chamber (24 °C, 55% relative humidity, 14-h light/10-h dark period). *Arabidopsis thaliana* plants were cultivated under short day conditions in a controlled environment chamber (21 °C, 55% relative humidity, 10-h light/14-h dark period). The following genotypes were used: *A. thaliana* Col-0 (wild type), the myrosinase deficient *A. thaliana tgg1×tgg2* (*tgg*) double knockout mutant (Barth and Jander, 2006), and the *A. thaliana myb28×myb29* (*myb*) double knockout mutant which does not produce aliphatic glucosinolates (Sønderby et al., 2007).

*Phyllotreta armoraciae* was reared on potted three- to four-week old *B. juncea* or *B. rapa* plants in a controlled environment chamber (24 °C, 60% relative humidity, 14-h light/10-h dark period). Adult beetles were provided with new plants every week and plants with eggs were kept separately for larval development. After three weeks, any remaining plant material was removed and the soil containing pupae was kept in plastic containers (9 L volume, Lock&Lock). Newly emerged adults were collected every two to three days. Unless stated otherwise, experiments were performed with newly emerged beetles that had been reared on *B. juncea* plants.

### Sequestration experiments

To analyze whether plant myrosinase activity influences the sequestration of glucosinolates in *P. armoraciae* beetles, we performed sequestration experiments with the myrosinase-deficient *Arabidopsis tgg* mutant and the corresponding wild type Col-0.

In *Experiment 1*, we fed newly emerged beetles for one day with detached leaves of *Arabidopsis* wild type or *tgg* plants (n = 28 per genotype, two beetles per replicate). On the next day, the remaining leaves were weighed, frozen in liquid nitrogen, and stored at −20 °C until they were freeze dried. To allow for metabolism of ingested aliphatic glucosinolates, we fed the beetles one additional day on *Arabidopsis myb* leaves before beetles were weighed, frozen in liquid nitrogen, and stored at −20 °C until extraction. The extraction and analysis of glucosinolates by high performance liquid chromatography coupled with diode array detection (HPLC-DAD) was performed as described in Beran et al. (2014). Adult *P. armoraciae* beetles convert ingested 4-methylsulfinylbutyl (4MSOB) glucosinolate, the major aliphatic glucosinolate in *Arabidopsis* wild type and *tgg* plants, into 4-methylthiobutyl (4MTB) glucosinolate (Yang et al., 2020). Therefore, we summed up the concentrations of 4MSOB and 4MTB glucosinolate in each beetle sample and expressed this concentration relative to that of both glucosinolates in the corresponding leaf sample, which was set to 100%. To confirm that beetles feed equally on both *Arabidopsis* lines, we quantified the beetle feeding damage (2 beetles per leaf disc with 16 mm diameter) over one day using the software Fiji (Schindelin et al., 2012) (n = 8 per genotype).

In *Experiment 2*, we fed newly emerged beetles with *Arabidopsis* wild type, *tgg*, or *myb* leaves for one day (n = 5, with 5 beetles per replicate). Afterwards, feces were collected in 50 μL ultrapure water containing 0.1% (v/v) formic acid, mixed with 50 μL pure methanol and stored at −20 °C. Beetles were frozen in liquid nitrogen and stored at −20 °C until extraction. Remaining leaves were weighed, frozen in liquid nitrogen and freeze-dried. Feces samples were homogenized for 2 min at 25 Hz in a TissueLyzerII (Qiagen) using metal beads. Beetles were homogenized in 500 μL 50% (v/v) methanol using plastic pestles. Freeze-dried leaves were homogenized to powder as described for feces samples and extracted with 800 μL of 50% (v/v) methanol. Samples were centrifuged at 4 °C for 10 min at 16,000 × g and supernatants were stored at −20 °C until analysis by liquid chromatography coupled with tandem mass spectrometry (LC-MS/MS). Identification and quantification of 4MSOB glucosinolate and 4MSOB glucosinolate-derived metabolites in samples was performed as described in Beran et al. (2018). After quantification, we subtracted the average amounts of each metabolite in bodies or feces of *myb*-fed control beetles (background control) from those detected in wild type- or *tgg*-fed beetles.

In *Experiment 3*, we fed newly emerged beetles for one day with *Arabidopsis* wild type or *tgg* leaves that were each spiked with 300 nmol allyl glucosinolate (Carl Roth) as described in Schramm et al. (2012). This experiment was performed with adult beetles that had been reared on *B. rapa* plants, which do not produce allyl glucosinolate (Beran et al., 2018). The glucosinolate profile of unfed beetles reared on *B. rapa* was analyzed as described in Beran et al. (2014) (n = 20, with 5 beetles per replicate) to confirm that allyl glucosinolate is not present in beetles. To prevent leaf wilting, we placed the petiole in a reaction tube filled with water. Newly emerged beetles were fed with spiked *Arabidopsis* leaves for one day (n = 10 per genotype, 5 beetles per replicate) and allyl glucosinolate-spiked leaves without beetles were kept under the same conditions as a recovery control (n = 8-10 per genotype). All leaf samples were frozen in liquid nitrogen, freeze-dried, and homogenized with metal beads (2.4 mm diameter, Askubal) for 2 min at 25 Hz in a TissueLyzerII (Qiagen). Beetles were weighed and frozen in liquid nitrogen. Feces were collected in 100 μL ultrapure water and mixed with 100 μL pure methanol. Leaf and beetle samples were homogenized in 1 mL and 800 μL 50% (v/v) methanol, respectively. After centrifugation for 10 min at 16,000 × g, supernatants were collected. Feces samples were homogenized as described for leaves and centrifuged at 4 °C or 10 min at 16,000 × g. The supernatant was collected, the solvent evaporated using nitrogen and extracts were re-dissolved in 80 μL 50% methanol. Samples were stored at −20 °C until LC-MS/MS as described in Malka et al. (2016) using a modified elution gradient. The gradient consisted of formic acid (0.2%) in water (solvent A) and acetonitrile (solvent B) and was carried out as follows: 1.5% (v/v) B (1 min), 1.5–5% (v/v) B (5 min), 5–7% (v/v) B (2 min), 7–12.6% (v/v) B (4 min), 12.6–100% (v/v) B (0.1 min), 100% (v/v) B (0.9 min), 100 to 1.5% (v/v) B (0.1 min), and 1.5% (v/v) B (3.85 min). Allyl glucosinolate was quantified using an external calibration curve. We recovered 97.8 ± 6.5% and 109.5 ± 4.4% (mean ± SD) of the spiked glucosinolate from undamaged (control) wild type and *tgg* leaves, respectively, showing that only small amounts of spiked allyl glucosinolate were metabolized in *Arabidopsis* leaves under our assay conditions. To determine how much allyl glucosinolate beetles had ingested, we subtracted the allyl glucosinolate amount detected in each fed leaf from the average allyl glucosinolate amounts recovered from corresponding unfed control leaves. The amounts of allyl glucosinolate that were recovered in beetles and feces were expressed relative to the total ingested amount, which was set to 100%.

In *Experiment 4*, we fed newly emerged beetles with allyl glucosinolate-spiked *Arabidopsis* wild type or *tgg* leaves (prepared as described in *Experiment 3*) for one day and simultaneously collected the headspace on Porapak-Q™ volatile collection traps (25 mg; ARS, Inc) (n = 6–7 per genotype, 8 beetles per replicate). Leaves without beetles served as controls (n = 4 per genotype). The volatile collection and sample analysis by gas chromatography mass spectrometry (GC-MS) was performed as previously described in Sporer et al. (2020). Allyl isothiocyanate was quantified in headspace samples using an external calibration curve prepared from an authentic standard (Sigma-Aldrich). The glucosinolate amount per fed beetle was determined as described in *Experiment 1*.

### Performance experiment

To investigate whether the hydrolysis of ingested glucosinolates has a negative influence on beetle performance, we compared the fresh weight and energy reserves of beetles that had fed for ten days on *Arabidopsis* wild type or *tgg* leaf discs. We separated newly emerged beetles into males and females and assigned them randomly to one of the two *Arabidopsis* genotypes (n = 10 females per genotype, n = 8-9 males per genotype). Each beetle was provided with a new leaf cut from an undamaged *Arabidopsis* plant every day for ten consecutive days. After ten days feeding, beetles were weighed, frozen in liquid nitrogen, and stored at −20 °C until analysis of energy reserves. The contents of soluble protein, total lipids, glycogen and soluble carbohydrates in individual beetles were determined as described in Foray et al. (2012) with minor modifications. Instead of a 96-well borosilicate microplate, we used a 96-well quartz glass microplate (Hellma Analytics) that was covered with MicroAmp clear adhesive film (Applied Biosystems). The plate was heated using a ThermoMixer (Eppendorf) and for measurements we used a Tecan Infinite 200 Reader (Tecan). As a control, we quantified the levels of soluble protein, amino acids, and sugars in rosette leaves of *Arabidopsis* wild type and *tgg* mutant plants (for details refer to Supplementary Methods).

### Short-term feeding experiment

In a short-term feeding experiment, we allowed newly emerged beetles to feed for 1 min on wild type or *tgg* leaves. After 5 min, beetles were dissected into gut and remaining body (without head; n = 3, 3 beetles per replicate). Non-fed beetles were used as background control (n = 2-3, 3 beetles per replicate). Dissected guts were washed twice in phosphate-buffered saline (PBS) pH 7.4 (Bio-Rad) before sampling. Samples were homogenized in 500 μL 80% methanol containing 0.4 μM 4-hydroxybenzyl glucosinolate as internal standard using plastic pestles and stored at −20 °C until extraction and analysis by LC-MS/MS as described above in *Experiment 2*. We quantified 4MSOB glucosinolate in each sample using an external standard curve and expressed the glucosinolate distribution in the gut and rest of the body relative to the total amount detected in both samples (set to 100%).

### Myrosinase inhibition assays

To determine whether *P. armoraciae* can inhibit ingested myrosinase activity, we performed myrosinase activity assays with gut content extracts of adult beetles. The gut content of adults was collected as follows: dissected guts were washed in extraction buffer (20 mM 2-(N-morpholino)ethanesulfonic acid (MES), pH 5.2) containing protease inhibitors (cOmplete, EDTA-free) and cut open longitudinally to collect the gut content in 2.5 μL of extraction buffer. For each sample, gut contents from 20 beetles were pooled, frozen in liquid nitrogen and stored at −20 °C until extraction (n = 4). Samples were homogenized with metal beads for 2 min at 25 Hz in a TissueLyzer II, centrifuged at 4 °C for 10 min at 16,000 × *g* and the supernatant split into two subsamples of which one was boiled for 5 min at 99 °C.

Assays (50 μL total volume) consisted of 0.1 mM ascorbic acid (Fluka, Buchs, Switzerland), 0.2 mM 4MSOB glucosinolate (substrate), 0.5 ng/μL partially purified myrosinase from *Sinapis alba* (Sigma-Aldrich, details of protein purification are described in the Supplementary Methods) and a) gut content extract (corresponding to four beetles), b) boiled gut content extract, or c) extraction buffer (control). Assays with gut extracts but without myrosinase and assays containing only substrate were used as additional controls. Assays were incubated for 15 min at 30 °C, the reaction was stopped by 5 min boiling at 99 °C and extracted with 100 μL 80% methanol containing 0.2 mM 4-hydroxybenzyl glucosinolate as an internal standard. Myrosinase activity was determined by quantifying the remaining 4MSOB glucosinolate amount in each assay as described in *Experiment 1*.

### Detection of myrosinase enzyme and activity in beetle feces

To determine whether *P. armoraciae* can degrade ingested plant myrosinase enzymes, we collected feces of 90 adults that had fed on *Arabidopsis* wild type leaves for one day in a total volume of 1 mL 20 mM MES buffer pH 6.5 containing protease inhibitors (cOmplete, EDTA-free, Roche). After homogenization with metal beads for 3 min at 25 Hz in a TissueLyzer II and centrifugation at 4 °C for 10 min at 16,000 × *g*, we precipitated soluble proteins in the supernatant using trichloroacetic acid and washed the pellet with acetone. The protein pellet was dissolved in Laemmli buffer (Bio-Rad), boiled for 15 min at 95 °C, and separated on a 12.5% Criterion Tris-HCl precast gel (Bio-Rad). Protein bands were stained with colloidal Coomassie G250 (Carl Roth), excised from the gel, and digested with porcine trypsin (Promega) as described in Shevchenko et al. (2006). Samples were re-dissolved in 30 μL 1% (v/v) formic acid and 2 μL were analyzed by nano-UPLC-MS^E^ analysis as described in Vassão et al. (2018). Data were acquired using data-independent acquisition, referred to as enhanced MS^E^. MS data were collected using MassLynx v4.1 software (Waters).

The processing of nano-UPLC-MS^E^ data and protein identification was performed as follows: the acquired continuum of LC-MS^E^ data were processed using the ProteinLynx Global Server (PLGS) version 2.5.2 (Waters) to generate product ion spectra for database searching according to the ion accounting algorithm described in Li et al. (2009). Processed data were searched against a reference sequence (Refseq) database containing *Arabidopsis thaliana* sequences (40785 sequences, downloaded from the Identical Protein Groups database at the National Center for Biotechnology Information (NCBI, https://www.ncbi.nlm.nih.gov/ipg on February 28, 2020) combined with a subdatabase containing common contaminants (O’Leary et al., 2016). Database searching was performed at a false discovery rate (FDR) of 2% with the following parameters: minimum numbers of fragments per peptide (3), peptides per protein (1), fragments per protein (7), and maximum number of missed tryptic cleavage sites (1). Searches were restricted to tryptic peptides with a fixed carbamidomethylation of cysteine residues along with variable oxidation of methionine. Proteins were classified according to the algorithm described for PAnalyzer software (Prieto et al., 2012) and divided into four groups: conclusive, indistinguishable, ambiguous, and non-conclusive. Conclusive and indistinguishable hits were considered as confident matches.

To determine whether *P. armoraciae* excretes active myrosinase enzyme, we analyzed myrosinase activity in feces homogenates and compared this activity with the corresponding ingested myrosinase activity in leaves. We used newly emerged *P. armoraciae* beetles that had been reared on *B. rapa*, and fed them for one day with *myb* leaves containing myrosinase activity but no 4MSOB glucosinolate (n = 6, with 6 beetles per replicate). Leaves were weighed before and after feeding to determine the ingested plant fresh weight. Leaves were supplied with water during the experiment and the average proportional weight gain of intact leaves was used to correct the initial leaf weight (n = 16). Feces from each replicate were collected in 130 μl extraction buffer (20 mM MES buffer, pH 6.5, containing protease inhibitors (cOmplete, EDTA-free)). Feces and fed leaves were frozen in liquid nitrogen and stored at −80 °C until extraction. Feces samples were homogenized with metal beads at 25 Hz for 2 min in a TissueLyzer II. The corresponding frozen leaf samples were homogenized with metal beads at 25 Hz for 2 min in a pre-cooled sample holder to prevent thawing. For each replicate, we calculated the ingested fresh weight and extracted the corresponding amount of homogenized plant tissue in the same buffer volume used for feces extraction. After centrifugation at 4 °C for 10 min at 16,000 × *g*, the supernatant was directly used for myrosinase activity assays. Assays consisted of 25 μL extraction buffer, 5 μL of an aqueous 11 mM 4MSOB glucosinolate solution, and a) 25 μL feces homogenate or b) 25 μL leaf extract. Assays containing only extraction buffer and substrate served as background control (n = 3). To test whether feces extracts have an inhibitory effect on plant myrosinase activity, we additionally performed assays in which we mixed 25 μL of feces homogenate with 25 μL of corresponding leaf extract and glucosinolate substrate. Except for three assays with combined feces and leaf extracts, all assays were performed with two technical replicates. Assays were incubated for 30 min at 30 °C, stopped by boiling for 5 min at 95 °C, and 50 μL of 60% (v/v) methanol were added. After the activity assay, samples containing feces homogenates were centrifuged at 4 °C for 10 min at 16,000 × g, the supernatant collected and final samples were stored at −20 °C until LC-MS/MS analysis. 4MSOB glucosinolate was quantified as described in *Experiment 2* and myrosinase activity was expressed as nmol 4MSOB hydrolyzed per minute and mg (ingested) plant fresh weight.

### Statistical analyses

Statistical analyses were performed in R3.3.1 (R Core Team, 2018) or in SigmaPlot 11.0 (Systat Software). Details of statistical analyses performed for each dataset are summarized in Supplementary Table 1.

## RESULTS

### Plant myrosinase activity influences glucosinolate sequestration

We examined the influence of plant myrosinase activity on glucosinolate sequestration by feeding *P. armoraciae* adults with *Arabidopsis* leaves with (wild type) or without (*tgg*) myrosinase activity (*Experiment 1*). Because *P. armoraciae* converts sequestered 4MSOB glucosinolate into 4MTB glucosinolate (Yang et al. 2020), we quantified the levels of both 4MSOB and 4MTB glucosinolate in beetles relative to those in the food plant and found that *tgg*-fed beetles accumulated twofold higher levels of glucosinolates than wild type-fed beetles (Mann-Whitney rank sum test, *U* = 482.000, *p* < 0.001; Figure 1A). Since beetle feeding rates and glucosinolate levels did not differ between treatments (feeding rate: Student’s *t*-test, *t* = 0.592, *p* = 0.564; plant glucosinolates level: Mann-Whitney rank sum test, *U* = 390.000, *p* = 0.980), our results demonstrate a negative impact of plant myrosinase activity on glucosinolate sequestration in *P. armoraciae*.

**Figure 1:**
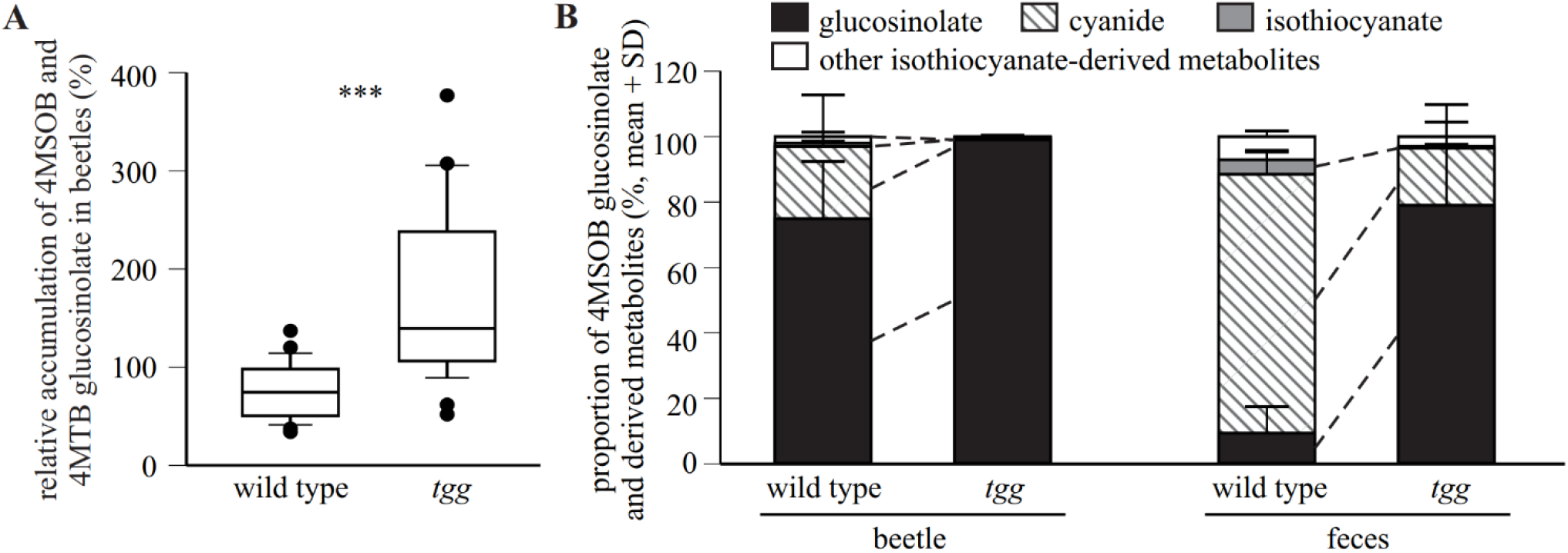
Plant myrosinase activity negatively influences the sequestration of 4MSOB glucosinolate in *P. armoraciae*. (A) Accumulation of 4-methylsulfinylbutyl (4MSOB) and 4-methylthiobutyl (4MTB) glucosinolate in *P. armoraciae* adults relative to the concentration in fed *Arabidopsis* wild type and myrosinase-deficient *tgg* leaves (n = 28). Glucosinolates were quantified after conversion to desulfo-glucosinolates by HPLC-DAD. The glucosinolate concentration in the plant was set to 100%. *** *p* < 0.001. (B) Relative composition of 4MSOB glucosinolate and hydrolysis products in bodies and feces of wild type- or *tgg*-fed adults (n = 5). Glucosinolates and hydrolysis products were extracted with 50% methanol and analyzed by LC-MS/MS. Detected amounts of metabolites were expressed relative to the total amounts of all detected metabolites in beetles or feces (set to 100%). Dashed lines indicate significant differences between samples (*p* < 0.05). Statistical results are shown in Supplementary Table 1. 4MSOB cyanide corresponds to the nitrile hydrolysis product of 4MSOB glucosinolate. Other isothiocyanate-derived metabolites comprise 4MSOB isothiocyanate-glutathione conjugate, 4MSOB isothiocyanate-cysteinylglycine conjugate, 4MSOB isothiocyanate-cysteine conjugate, 2- (4-(methylsulfinyl)butylamino)-4,5dihydrothiazole-carboxylic acid, 4MSOB amine, and 4MSOB acetamide.

To determine whether the lower 4MSOB glucosinolate accumulation rate in wild type-compared to *tgg*-fed beetles is due to hydrolysis, we quantified the levels of 4MSOB glucosinolate and known hydrolysis products in bodies and feces of wild type- and *tgg*-fed beetles by LC-MS/MS (*Experiment 2*). We found that the detected metabolite levels differed greatly between replicates (Supplementary Table 2), despite similar 4MSOB glucosinolate concentrations in wild type- and *tgg-*leaves (Student’s *t*-test, *t* = 0.371, *p* = 0.720). We thus compared the relative composition of metabolites between treatments and found significantly higher percentages of hydrolysis products in body and feces samples of wild type-fed beetles than in the corresponding samples of *tgg*-fed beetles (Figure 1B; results of statistical analyses are summarized in Supplementary Table 1). 4MSOB cyanide represented the dominant hydrolysis product in body and feces samples whereas only traces of free and metabolized isothiocyanates were detected (Supplementary Table 2, Figure 1B). Overall, glucosinolate hydrolysis products accounted for 28% and 2.2% of the total detected metabolites in wild type- and *tgg*-fed beetles, respectively (bodies and feces). Together, our results show that a fraction of ingested 4MSOB glucosinolate was hydrolyzed by the plant myrosinase, but that glucosinolate hydrolysis also occurred independently of plant myrosinase activity.

To directly quantify the impact of plant myrosinase activity on the metabolic fate of ingested glucosinolates in *P. armoraciae*, we performed a feeding experiment with wild type- and *tgg*-leaves that were spiked with allyl glucosinolate, the major glucosinolate of the natural host plant of *P. armoraciae* (*Experiment 3*). This experiment was performed with adult beetles that were reared on *B. rapa* plants and, therefore, do not contain allyl glucosinolate (Supplementary Table 3). *Tgg-fed* beetles accumulated a significantly higher proportion of ingested allyl glucosinolate than wild type-fed beetles (Table 1). Plant myrosinase activity explained the fate of 9% of the ingested allyl glucosinolate; however, the metabolic fate of 47% and 56% of the ingested allyl glucosinolate in *tgg-* and wild type-fed beetles remained unknown, respectively (Table 1).

**Table 1:**
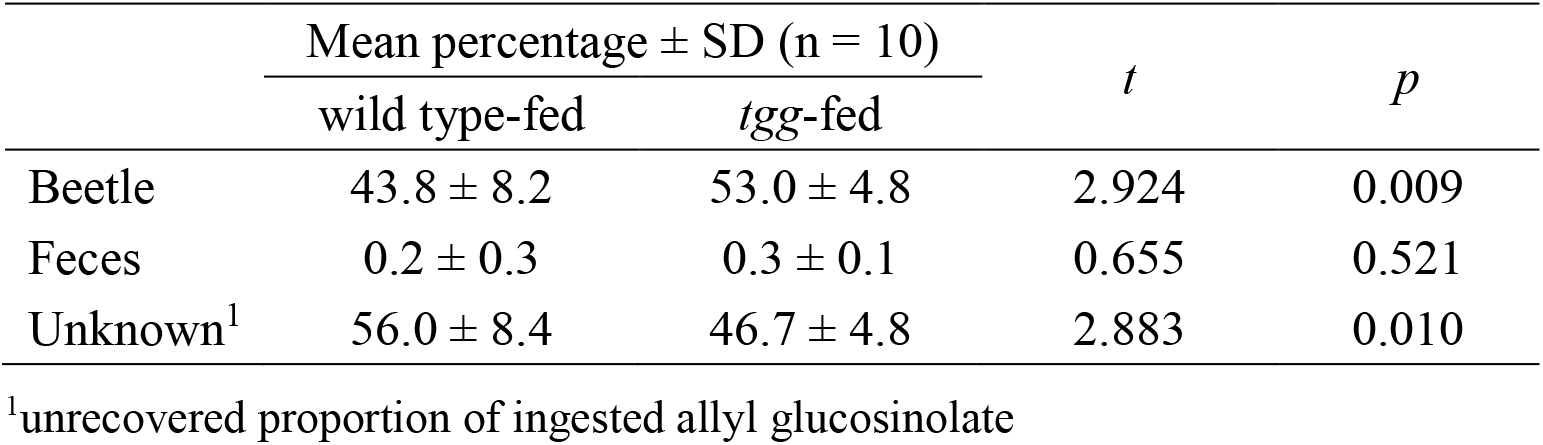
Recovery of ingested allyl glucosinolate in *P. armoraciae*.

| | Mean percentage ± SD (n = 10) | | *t* | *p* |
| --- | --- | --- | --- | --- |
|  | wild type-fed | *tgg*-fed |  |  |
| Beetle | 43.8 ± 8.2 | 53.0 ± 4.8 | 2.924 | 0.009 |
| Feces | 0.2 ± 0.3 | 0.3 ± 0.1 | 0.655 | 0.521 |
| Unknown<sup>1</sup> | 56.0 ± 8.4 | 46.7 ± 4.8 | 2.883 | 0.010 |

<sup>1</sup>unrecovered proportion of ingested allyl glucosinolate

To confirm that ingested allyl glucosinolate is hydrolyzed, we quantified the emission of the volatile glucosinolate hydrolysis product allyl isothiocyanate during beetle feeding (*Experiment 4*). We detected significantly higher amounts of allyl isothiocyanate in the headspace samples of wild type-fed beetles as compared to *tgg*-fed beetles, while the levels of sequestered allyl glucosinolate were not influenced by the food plant (Figure 2; results of statistical analyses are summarized in Supplementary Table 1).

**Figure 2:**
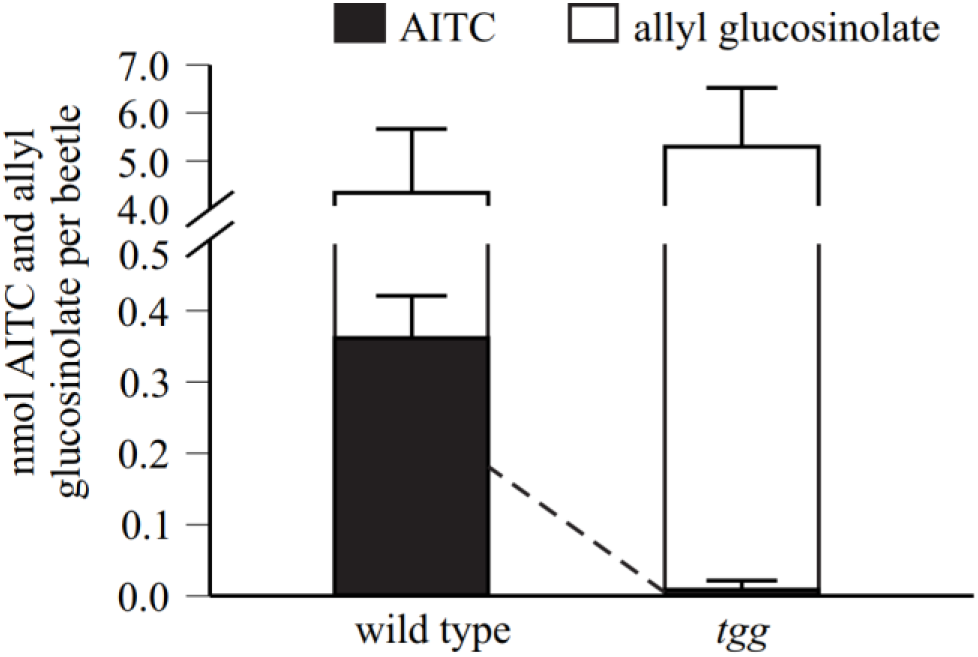
Allyl glucosinolate is hydrolyzed during *P. armoraciae* feeding on *Arabidopsis* wild type leaves. Adult beetles were fed with allyl glucosinolate-spiked *Arabidopsis* leaves with (wild type) and without (*tgg*) myrosinase activity for one day. Headspace volatiles were collected on Porapaq-Q^TM^ adsorbent, eluted with hexane, and allyl isothiocyanate was quantified using gas chromatography-mass spectrometry (*m/z* 99). Detected amounts of allyl isothiocyanate were corrected by subtracting the background emission detected in volatile collections performed without beetles, which served as control. Allyl glucosinolate was quantified after conversion to desulfo-glucosinolates by HPLC-DAD. The dashed line indicates significant differences between samples (*p* < 0.001; n = 6-7). Results of statistical analyses are provided in Supplementary Table 1.

### Plant myrosinase activity does not affect beetle weight and energy reserves

To determine whether the hydrolysis of ingested glucosinolates by plant myrosinase influences beetle performance, we compared the beetle fresh weight and the levels of soluble proteins, carbohydrates, glycogen and lipids in beetles that were fed with *Arabidopsis* leaves with and without myrosinase activity for ten days. The food plant had no influence on the beetle weight or energy reserves, indicating that *P. armoraciae* adults can tolerate the partial hydrolysis of ingested glucosinolates (Supplementary Table 4). As a control, we compared the nutritional value of wild type and *tgg* leaves and also found no differences regarding the total levels of soluble protein, free amino acids, soluble sugars and glucosinolates between these genotypes (Supplementary Table 5).

### *P. armoraciae* rapidly sequesters glucosinolates and suppresses plant myrosinase activity in the gut

The rapid absorption of ingested glucosinolates across the gut epithelium represents one possible mechanism to prevent hydrolysis in the gut lumen. We analyzed the distribution of ingested glucosinolates in *P. armoraciae* beetles 5 min after feeding on wild type and *tgg* leaves and recovered more than 80% of the total detected 4MSOB glucosinolate from the body (without gut). The relative distribution of detected glucosinolates did not differ between wild type- and *tgg*-fed beetles (Student’s *t*-test, *t* = 0.075, *p* = 0.944, Figure 3A). To determine whether glucosinolate sequestration is facilitated by suppression of plant myrosinase activity in the gut lumen of *P. armoraciae*, we analyzed the influence of beetle gut content extracts on activity of partially purified *S. alba* myrosinase in *in vitro* assays. Compared to control assays, gut content extracts significantly reduced myrosinase activity by up to 50% (ANOVA, *F* = 85.639,*p* < 0.001). Boiled gut content extracts reduced myrosinase activity significantly less than untreated extracts (Figure 3B).

**Figure 3:**
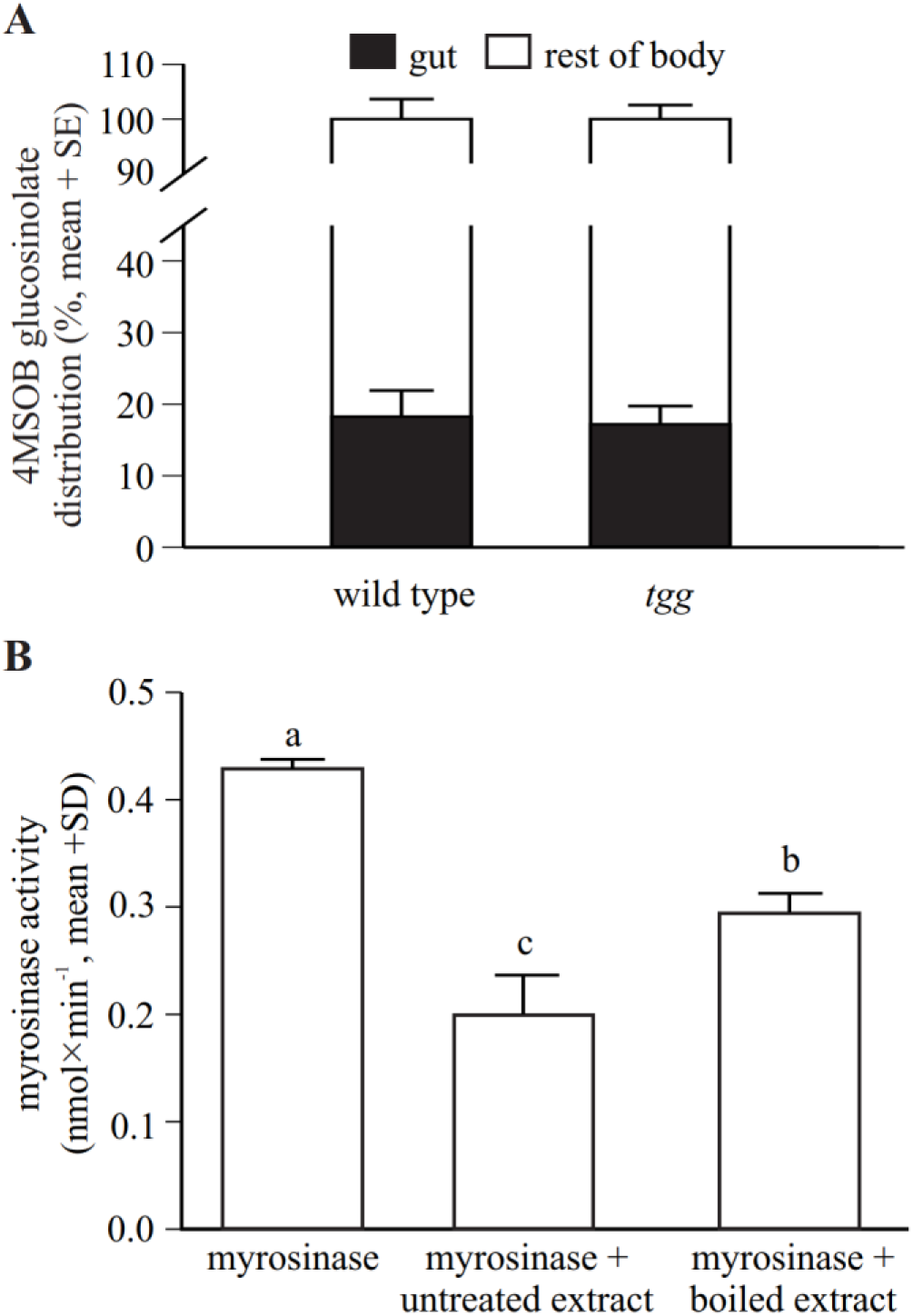
*P. armoraciae* rapidly sequesters ingested glucosinolates and can suppress plant myrosinase activity. (A) Beetles were allowed to feed for 1 min on *Arabidopsis* leaves with (wild type) or without (*tgg*) myrosinase activity and were dissected into gut and rest of body 5 min later (n = 3). Beetles were extracted with 80% methanol and 4MSOB glucosinolate was quantified by LC-MS/MS. The distribution of 4MSOB glucosinolate in gut and rest of body is expressed relative to the total amount detected in both samples (set to 100%). (B) Partially purified *Sinapis alba* myrosinase was affected by the supplementation of untreated and boiled gut content extracts in *in vitro* assays (n = 4). Myrosinase activity was determined by quantifying the 4MSOB glucosinolate substrate in each assay after conversion to desulfo-glucosinolate and analysis by HPLC-DAD. Assays without myrosinase served as background controls and activities were subtracted from the corresponding samples. Different letters indicate significant differences, *p* < 0.001.

### *P. armoraciae* excretes inactive myrosinase enzyme

To investigate whether the *Arabidopsis* myrosinases TGG1 and TGG2 are degraded in the gut of *P. armoraciae*, we analyzed the fecal proteome by nano-UPLC-MS^E^. We detected a total of 14 peptides derived from the *Arabidopsis* myrosinase TGG1 in two protein bands between 55 and 70 kDa that covered 34% of the TGG1 amino acid sequence (Figure 4A, B; Supplementary Table 6). The molecular weight range in which we detected TGG1 peptides corresponds approximately to the predicted (61.1 kDa) and apparent (75 kDa) molecular weight of TGG1 (Zhou et al., 2012). TGG2-specific peptides were not detected in beetle feces with our approach (Supplementary Data 1).

**Figure 4:**
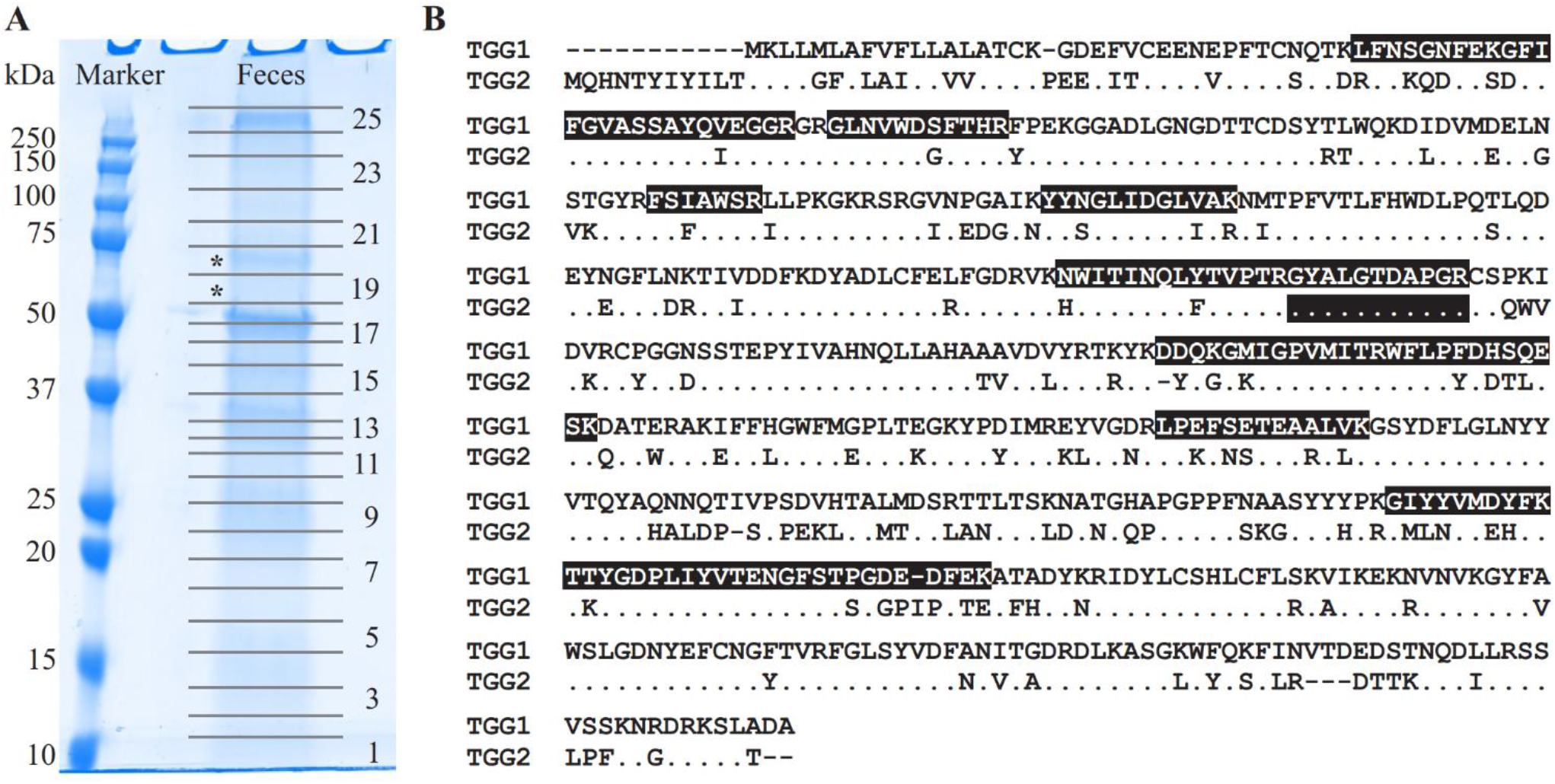
Detection of ingested *Arabidopsis* myrosinase in the feces of *P. armoraciae*. (A) One dimensional SDS/PAGE gel of crude feces protein extract. Numbers indicate samples excised for proteomic analysis by nano-UPLC-MS^*E*^ (only odd numbers are shown). TGG1-derived peptides were detected in bands marked with asterisks. (B) Amino acid sequence alignment of the *Arabidopsis* myrosinases TGG1 (AT5G26000.1) and TGG2 (AT5G25980.2). Identical amino acids in the TGG2-sequence are represented by a dot. Peptides highlighted with black background were detected by nano-UPLC-MS^*E*^. Only one detected peptide matched both TGG1 and TGG2.

Since the proteomic analysis indicates that *P. armoraciae* excretes intact myrosinase, we compared the levels of ingested myrosinase activity with those that were excreted. Myrosinase activity detected in feces corresponded to less than 4% of the ingested activity (Figure 5, *t* = 10.449, *p* < 0.005). We additionally spiked plant myrosinase extracts into feces homogenates, but observed similar activity as in control assays (Figure 5, *t* = 0.158, *p* = 1.000).

**Figure 5:**
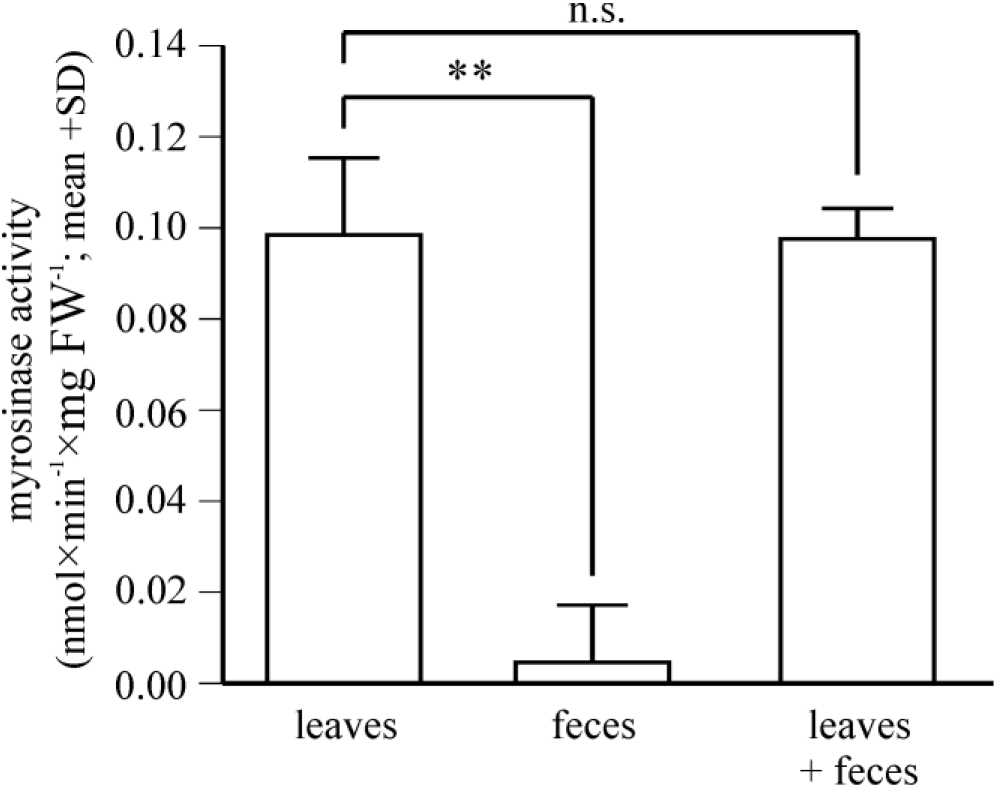
*Arabidopsis* myrosinase activity is strongly reduced after digestion. Enzymatic levels of myrosinase activity in *Arabidopsis myb* leaves were determined in non-ingested leaf material (leaves) and after beetle digestion (feces) of equivalent amounts. Samples were spiked with 4MSOB glucosinolate and the reaction was stopped by heat inactivation after 30 min. Remaining glucosinolate substrate was quantified by LC-MS/MS analysis. Co-incubation of non-ingested myrosinases with *P. armoraciae* feces homogenate did not affect myrosinase activity levels. **, *p* < 0.005; n.s., *p* < 0.05; n = 6. Statistical results are shown in Supplementary Table 1.

## DISCUSSION

The enzymatic activity of defensive plant β-glucosidases is a major barrier to the sequestration of plant glucosides by herbivorous insects (Morant et al., 2008; Pentzold et al., 2014b). Here, we investigated the impact of plant myrosinase activity on the sequestration of glucosinolates in *P. armoraciae* and explored possible mechanisms that enable adult beetles to suppress glucosinolate hydrolysis during feeding and digestion. We demonstrate a negative influence of plant myrosinase activity on sequestration and confirmed that a fraction of ingested glucosinolates is hydrolyzed by plant myrosinases. We found two mechanisms that can reduce the glucosinolate hydrolysis rate in the gut: the rapid absorption of ingested glucosinolates across the gut epithelium and the suppression of plant myrosinase activity in the gut lumen.

Compared to previous studies with *P. chrysocephala* adults (Beran et al., 2018; Ahn et al., 2019), plant myrosinase activity had less impact on the metabolic fate of ingested glucosinolates in *P. armoraciae* adults. In quantitative studies, plant myrosinases hydrolyzed approximately 75% of the total ingested glucosinolates in *P. chrysocephala*, whereas only about 10% of ingested allyl glucosinolate were hydrolyzed in *P. armoraciae*. Although *P. chrysocephala* and *P. armoraciae* belong to different genera, they have similar body sizes and feeding modes and therefore cause comparable feeding damage. Our results thus indicate that *P. armoraciae* adults are better adapted to overcome plant myrosinase activity than *P. chrysocephala* adults.

Plant myrosinases hydrolyzed a fraction of ingested glucosinolates in *P. armoraciae* and thus negatively influenced glucosinolate sequestration (Figure 1 and 2, Table 1). Although the sequestration rates of 4MSOB glucosinolate and allyl glucosinolate cannot be compared directly because different methods were used for quantification, our results indicate a stronger influence of plant myrosinase activity on 4MSOB glucosinolate than on allyl glucosinolate. Biochemical studies with *Arabidopsis* myrosinases TGG1 and TGG2 revealed similar activities of both enzymes towards these two glucosinolates (Zhou et al., 2012), making it unlikely that the substrate preferences of *Arabidopsis* myrosinases affected sequestration. A similar glucosinolate-dependent effect of plant myrosinase activity on sequestration was also observed in *P. armoraciae* larvae (Sporer et al., 2020). Larvae sequestered allyl glucosinolate from *B. juncea* leaves but almost no glucosinolates from *Arabidopsis* wild type leaves, despite similar levels of soluble myrosinase activity in both plant species. Thus, the metabolic fate of ingested allyl glucosinolate in *P. armoraciae* appears to be less affected by plant myrosinase activity than that of *Arabidopsis* glucosinolates. Since allyl glucosinolate represents the dominant glucosinolate in horseradish, *P. armoraciae* might have developed specific mechanisms to avoid the hydrolysis of the characteristic glucosinolate of its natural host plant (Li and Kushad, 2004).

Glucosinolate hydrolysis products, in particular isothiocyanates, are well-known to have a negative impact on insect growth and development by interfering with nutrition (Agrawal and Kurashige, 2003; Jeschke et al., 2016b; Jeschke et al., 2017; Sun et al., 2019). Under our experimental conditions, the hydrolysis of ingested glucosinolates by plant myrosinases did not influence the weight or energy reserves of *P. armoraciae* adults. In addition, we observed no effect of plant myrosinase activity on developmental time, weight, and energy reserves of *P. armoraciae* larvae (for details refer to Supplementary Materials and Supplementary Table 9). The exposure to glucosinolate hydrolysis products could influence other fitness parameters such as female fecundity or egg hatching rate (Sun et al., 2019); however, our current results indicate that *P. armoraciae* can tolerate glucosinolate hydrolysis without a major impact on beetle performance.

Previous feeding experiments performed with generalist lepidopteran herbivores revealed 4MSOB isothiocyanate to be the major hydrolysis product of 4MSOB glucosinolate in *Arabidopsis* (Schramm et al., 2012; Jeschke et al., 2017). However, we detected much more 4MSOB cyanide in *P. armoraciae* bodies and feces relative to 4MSOB isothiocyanate and its derivatives (Figure 1B). There are several possible explanations for this unexpected result: 4MSOB isothiocyanate may not have been recovered completely with our extraction methods because it reacted with proteins or was metabolized by the beetle or associated gut microbes (Brown and Hampton, 2011; Jeschke et al., 2015; van den Bosch and Welte, 2017; Friedrichs et al., 2020; Shukla and Beran, 2020). Alternatively, *P. armoraciae* may be able to manipulate the outcome of glucosinolate hydrolysis and promote the formation of less toxic 4MSOB cyanide instead of 4MSOB isothiocyanate. Manipulation of glucosinolate hydrolysis occurs for example in larvae of the cabbage white butterfly, which express a so-called nitrile specifier protein in the gut (Wittstock et al., 2004). In fact, a redirection of glucosinolate hydrolysis towards less toxic nitriles could explain why plant myrosinase activity had no measurable effect on *P. armoraciae* performance. Further research is needed to understand the mechanism underlying the unusual composition of glucosinolate hydrolysis products in *P. armoraciae*.

How chewing insects that cause extensive tissue damage prevent the hydrolysis of ingested plant glucosides is currently not well understood. One proposed mechanism is a rapid absorption of plant glucosides across the gut epithelium that separates substrate and enzyme (Abdalsamee et al., 2014; Pentzold et al., 2014b; van Geem et al., 2014). Indeed, already after few minutes, we detected most ingested glucosinolates in the beetle body (without gut). This shows that glucosinolate uptake occurs rapidly in *P. armoraciae*, but it is unclear whether this prevents glucosinolate hydrolysis in the gut lumen. Moreover, we had expected to find less glucosinolates in the guts of wild type-fed beetles due to the presence of myrosinase activity, but we recovered similar proportions of glucosinolates in the guts of *tgg*- and wild type-fed beetles. However, we cannot rule out that glucosinolates detected in the gut were spatially separated from plant myrosinases, either in remaining intact plant tissue or in the gut epithelium.

Another explanation for the detection of glucosinolates in the gut of *P. armoraciae* is the inhibition of plant myrosinase activity in the gut. We found that gut content extracts reduced plant myrosinase activity by 30 to 50% in *in vitro* assays, with boiled extracts inhibiting myrosinase activity significantly less than untreated extracts. These results provide direct evidence for suppression of myrosinase activity in the gut of *P. armoraciae* and indicate that several factors contribute to myrosinase inhibition, of which some are sensitive to heat. Candidate agents that can modify plant myrosinase activity include ascorbic acid (cofactor of myrosinases), sulfate, sodium chloride and silver ions (Shikita et al., 1999; Andersson et al., 2009; Bhat and Vyas, 2019; Marcinkowska and Jelen, 2020). In addition, the gut pH can have a strong influence on the activity of ingested plant enzymes (Pentzold et al., 2014b). For example, the highly alkaline pH of the midgut lumen of burnet moth larvae drastically reduced cyanogenic β-glucosidase activity in *Lotus corniculatus* leaf macerates (Pentzold et al., 2014a). In contrast, the neutral pH of gut homogenates of glucosinolate-sequestering turnip sawfly larvae had only minor influence on ingested plant myrosinase activity (Abdalsamee et al., 2014). In *P. armoraciae*, gut homogenates showed an acidic pH (details are described in the Supplementary Material), which is unlikely to have a strong influence on plant myrosinase activity.

Myrosinase from *S. alba* and other defensive β-glucosidases were largely resistant to digestion in the larval gut of the generalist lepidopteran *Spodoptera littoralis* and thus retained most of the activity after digestion (Vassão et al., 2018). Our proteomic analysis of beetle feces also indicates that *Arabidopsis* myrosinase TGG1 resisted digestion in *P. armoraciae*, whereas TGG2 was not detected in feces (Figure 4, Supplementary Table 7). Because *TGG2* expression is restricted to phloem-associated cells (Barth and Jander, 2006), beetles likely ingested much less TGG2 than TGG1 by avoiding the leaf midrib and veins (personal observation). Despite the detection of TGG1 enzyme, we found almost no myrosinase activity in feces of *P. armoraciae*. We tested for the presence of myrosinase inhibitor(s) in feces homogenates but observed no suppression of spiked myrosinase activity under our assay conditions (Figure 5). Thus, we hypothesize that ingested TGG1 has been inactivated during gut passage in *P. armoraciae*. Previous studies with the turnip sawfly and the diamondback moth also indicated that plant myrosinases are not fully active in the gut (Abdalsamee et al., 2014; Sun et al., 2019). However, the underlying mechanism(s) of myrosinase inhibition in the gut of specialist herbivores including *P. armoraciae* remain to be determined.

*P. armoraciae* possesses endogenous myrosinase activity, which enables this specialist to exploit sequestered glucosinolates for defense against predators (Sporer et al. 2020). Our study provides first evidence that beetle myrosinases also contribute to glucosinolate metabolism in both adults (Figure 1B) and larvae (refer to Supplementary Materials, Supplementary Table 8 and Supplementary Figure 1). For example, *P. armoraciae* adults emitted the volatile hydrolysis product allyl isothiocyanate during feeding on myrosinase-deficient *tgg* leaves containing allyl glucosinolate. Volatile hydrolysis products derived from sequestered glucosinolates have also been detected in the headspace of the striped flea beetle, *Phyllotreta striolata*, which led to the initial discovery of a glucosinolate-myrosinase defense system in *Phyllotreta* flea beetles (Beran, 2011; Beran et al., 2014). To investigate the role of the beetle myrosinase in glucosinolate metabolism and defense in *P. armoraciae*, we currently perform experiments using beetles with suppressed myrosinase activity.

## CONCLUSION

Defensive plant β-glucosidases represent a major target of herbivore adaptation to two-component chemical defenses. Our study demonstrates that a specialist herbivore, *P. armoraciae*, can prevent the hydrolysis of glucosinolates using at least two strategies, the rapid absorption of the glucosinolates across the gut epithelium and the inhibition of plant myrosinase activity in the gut lumen. However, we also found that *P. armoraciae* can tolerate glucosinolate hydrolysis and is thus well-adapted to the chemical defense in its brassicaceous host plants.

## AUTHOR CONTRIBUTIONS

TS, JK, and FB designed experiments, TS, JK, YH and FB performed experiments, TS, JK, MR, NW and FB analyzed data, SGJ performed bioinformatic analyses, TS, JK, and FB wrote the manuscript.

## ACKNOWLEDGEMENTS

We thank the greenhouse team at the Max Planck Institute for Chemical Ecology for plant cultivation, Susanne Donnerhacke, Alexander Schilling, Fabian Seitz, and Leopold Wohlsperger for help with the rearing and experiments, Grit Kunert for help with statistical analyses, Daniel Veit and the workshop team for technical support, Caroline Müller and Helga Pankoke (University of Bielefeld) for help with the gut pH measurements, Sarah Wolf (Agroscope, Switzerland) for lending equipment for the energy budget analysis, and Felix Feistel for discussions.

## DATA AVAILABILITY STATEMENT

Source data of this study is available at the open access data repository of the Max Planck Society (Edmond) under https://dx.doi.org/10.17617/3.5b.

