## Supplementary Information for "A specialist flea beetle manipulates and tolerates the activated chemical defense in its host plant"

### *Supplementary Material*

#### SUPPLEMENTARY METHODS AND RESULTS

##### Chemical analyses of *Arabidopsis* wild type and *tgg* plants

For chemical analyses of *Arabidopsis* wild type and *tgg* plants, we harvested whole rosettes of six-week old plants (n = 10). Rosettes were frozen in liquid nitrogen, freeze-dried and homogenized to plant powder by shaking with metal beads (2.4 mm diameter, Askubal) for 2 min at 25 Hz in a TissueLyser II (Qiagen). Glucosinolates were extracted from 20 mg plant powder, converted to desulfo-glucosinolates and analyzed by HPLC-DAD as described in Beran et al. (2014). To determine the soluble protein content, 10 mg plant powder were extracted with 900 µL of 20 mM MES buffer pH 6.5 by shaking with metal beads for 2 min at 20 Hz in a TissueLyser II, followed by centrifugation at 4 °C for 10 min at 16,000 × g. Protein levels in supernatants were determined using the Bradford protein assay (Bio-Rad). Amino acids and sugars were analyzed by LC-MS/MS. Therefore, 15 mg plant powder were extracted with 1 mL of 80% MeOH by shaking and centrifugation as described above. The supernatant was diluted 1:10 with water containing a mix of $^{15}N/^{13}C$ labeled algal amino acids at a concentration of 10 µg × $mL^{-1}$ (Isotec). Amino acids were analyzed by LC-MS/MS on a Zorbax Eclipse C18-column (XDB-C18, 50 x 4.6 mm x 1.8 µm; Agilent, Santa Clara, CA, USA) (for details, refer to Crocoll et al. (2016)). Each amino acid was quantified relative to the peak area of its corresponding labeled amino acid, except for tryptophan (using phenylalanine and applying a response factor of 0.42) and asparagine (using aspartate and a response factor of 1.0). Soluble sugars were analyzed from the 1:10-diluted extract by LC-MS/MS on a hydrophilic interaction liquid chromatography (HILIC)-column (apHera-$NH_2$ Polymer; 15 × 4.6 mm, 5 µm; Supelco) as described in Madsen et al. (2015). Sugars were quantified using external standard curves prepared from authentic standards of glucose, fructose, sucrose (Sigma-Aldrich) and raffinose (Fluka).

##### Partial purification of myrosinase from *Sinapis alba*

We purified commercially available myrosinase enzyme (100 units (U) per g solid, isolated from *Sinapis alba* seeds; Sigma Aldrich) by fast protein liquid chromatography. The crude enzyme extract (ca. 25 U) was dissolved in 1 mL buffer (20 mM Tris·HCl, 0.15 M NaCl, pH 8, containing protease inhibitors (cOmplete, EDTA-free)) and subjected to size exclusion chromatography using a Superdex 200 10/300 GL column (GE Healthcare) as described in Beran et al. (2014). Fractions were tested for myrosinase activity as described below. Active fractions were pooled and desalted using Zeba Spin Desalting Columns (7 KDa MWCO 5 mL; Thermo Fisher Scientific) equilibrated with 20 mM Tris HCl pH 8 before anion exchange chromatography using a 1 mL ResourceQ column (GE Healthcare) as described in Beran et al. (2014). Fractions containing myrosinase activity were pooled and dialyzed overnight against 20 mM MES buffer pH 6.5 containing 20% (v/v) glycerol at 4 °C using a Slide-A-Lyzer Dialysis Cassette (10K MWCO, 3 mL, Thermo Fisher Scientific). After dialysis, protease inhibitors (cOmplete, EDTA-free, Roche) were added, the protein concentration determined using the Bradford protein assay (Bio-Rad) and the extract was stored at 4 °C.

Protein fractions were screened for myrosinase activity using a protocol modified from Travers-Martin et al. (2008). Assays consisted of 5 µL sample, 45 µL 20 mM MES buffer (pH 6.5) containing 2.78 mM allyl glucosinolate (Carl Roth) as substrate, and 50 µL assay reagent (20 mM MES buffer (pH 6.5) containing 57 U/ml glucose-oxidase (E.C.1.1.3.4, from *Aspergillus niger*, Serva), 5.6 U/ml peroxidase (E.C.1.11.1.7, from horseradish, Serva), 30.7 mM phenol (Sigma Aldrich) and 2.8 mM 4-aminoantipyrine (Sigma Aldrich)). Assays were incubated at room temperature for 30 min in transparent polystyrene 96-well microplates (Nunc, Thermo Fisher Scientific). Myrosinase activity was visually detected (pink assay color).

**pH measurements of *P. armoraciae* gut homogenates**

We dissected the midgut of adult *P. armoraciae* beetles collected from *B. juncea* rearing cages (n = 6). Dissected midguts (containing plant material) were homogenized individually in 40 µL deionized water and the pH of the obtained midgut homogenate was measured using an InLab Micro electrode (Mettler-Toledo, Schwerzenbach, Switzerland). The pH of midgut homogenates of *P. armoraciae* was 4.7 ± 0.2 (mean ± SD).

**Sequestration experiment with *P. armoraciae* larvae**

Previous experiments suggested that *P. armoraciae* larvae that feed on *Arabidopsis* wild type leaves are not able to prevent hydrolysis of ingested glucosinolates (Sporer et al., 2020). To investigate the influence of plant myrosinase activity on the metabolic fate of ingested glucosinolates in *P. armoraciae* larvae, we performed a feeding experiment with *Arabidopsis* wild type, *tgg* and *myb* plants, and quantified the levels of 4MSOB glucosinolate and its hydrolysis products in larvae and feces. The experimental set-up followed *Experiment 2* described in the main manuscript, except that the midribs of *Arabidopsis* leaves were removed using a scalpel to prevent larvae from mining. Feces, larvae and remaining leaves were sampled, extracted and analyzed by LC-MS/MS (for 4MSOB glucosinolate and derived metabolites) as described in *Experiment 2* (n = 5, with 5 larvae per replicate). The levels of 4MSOB glucosinolate in fed leaves did not differ significantly (Student's $t$-test, $t = 0.132$, $p = 0.898$). The detected amounts of 4MSOB glucosinolate and its hydrolysis products in larvae and feces are summarized in Supplementary Table 9.

In agreement with previous data, *tgg*-fed larvae sequestered higher levels of 4MSOB glucosinolate than wild type-fed larvae (Supplementary Table 9). Because the total levels of glucosinolate-derived metabolites differed greatly between replicates, we compared their relative composition between treatments (Supplementary Figure 1, results of statistical analyses are summarized in Supplementary Table 1). This comparison revealed a food plant-dependent composition of glucosinolate-derived metabolites in larval bodies. In wild type-fed larvae we detected predominantly 4MSOB cyanide, whereas 4MSOB glucosinolate, 4MSOB isothiocyanate and 4MSOB isothiocyanate-derived metabolites were more abundant in *tgg*-fed larvae (Supplementary Figure 1). Feces of wild type- and *tgg*-fed larvae contained similar percentages of mainly 4MSOB glucosinolate and 4MSOB cyanide (Supplementary Table 1). Overall, glucosinolate hydrolysis accounted for 95% and 76% of the total detected metabolites in wild type- and *tgg*-fed larvae, respectively (bodies and feces). These results indicate that *P. armoraciae* larvae have a much higher glucosinolate turnover rate compared to adults (Figure 1B), and that this turnover is largely independent of plant myrosinase activity.

**Performance experiment with *P. armoraciae* larvae**

We performed a long-term feeding experiment to investigate the influence of plant myrosinase activity on developmental time, fresh weight, and the energy budget of *P. armoraciae* larvae. Early second instar larvae were randomly assigned to either wild type or *tgg*, and were regularly provided with fresh leaves until they developed into prepupae. We recorded the developmental time and the final weight of each individual (n = 68-74), froze the prepupae in liquid nitrogen, and stored them at -20 °C until analysis of energy reserves (performed as described in the main manuscript; n = 11 per *Arabidopsis* genotype, with two individuals per replicate). The food plant had no influence on developmental time, fresh weight, and energy reserves of *P. armoraciae* larvae (Supplementary Table 9).

#### SUPPLEMENTARY TABLES

**Supplementary Table 1:** Methods and results of statistical analyses.

| Experiment | Comparison | Statistical method | Variable | Statistics | *p* |
|---|---|---|---|---|---|
| Experiment 1 | Relative 4MSOB and 4MTB glucosinolate accumulation in beetles (wild type vs. *tgg*) | Mann-Whitney rank sum test | Percentage[A] | *U* = 482.000 | < 0.001 |
| | Beetle feeding rate ($cm^2$) (wild type vs. *tgg*) | Student's *t*-test | Leaf area | *t* = 0.592 | 0.564 |
| | Plant 4MSOB and 4MTB glucosinolate levels (wild type vs. *tgg*) | Mann-Whitney rank sum test | Concentration | *U* = 390.000 | 0.980 |
| Experiment 2 | 4MSOB glucosinolate in beetles (wild type vs. *tgg*) | Student's *t*-test[B] | Percentage[A] | *t* = 3.931 | 0.016 |
| | 4MSOB-cyanide in beetles (wild type vs. *tgg*) | | | *t* = 3.963 | 0.016 |
| | 4MSOB isothiocyanate in beetles (wild type vs. *tgg*) | Mann-Whitney rank sum test[B] | | *U* = 0.000 | 0.032 |
| | Other 4MSOB isothiocyanate derived metabolites in beetles (wild type vs. *tgg*) | | | *U* = 0.000 | 0.032 |
| | 4MSOB glucosinolate in feces (wild type vs. *tgg*) | Student's *t*-test[B] | | *t* = 6.389 | 0.004 |
| | 4MSOB-cyanide in feces (wild type vs. *tgg*) | | | *t* = 6.686 | 0.004 |
| | 4MSOB isothiocyanate in feces (wild type vs. *tgg*) | | | *t* = 3.913 | 0.016 |
| | Other 4MSOB isothiocyanate derived metabolites in (wild type vs. *tgg*) | Mann-Whitney rank sum test[B] | | *U* = 5.000 | 0.604 |
| | 4MSOB glucosinolate and derived metabolites in beetles (wild type vs. *tgg*) | Student's *t*-test, Mann-Whitney rank sum test | Amount | see Supplementary Table 2 | |
| | 4MSOB glucosinolate and derived metabolites in beetle feces (wild type vs. *tgg*) | | | | |
| | Plant 4MSOB glucosinolate levels (wild type vs. *tgg*) | Student's *t*-test | Concentration | *t* = 0.371 | 0.720 |
| Experiment 3 | Recovery of ingested allyl glucosinolate in beetles and feces (wild type vs. *tgg*) | Student's *t*-test | Percentage[A] | see Table 1 | |
| Experiment 4 | Emitted allyl isothiocyanate per beetle (allyl glucosinolate-spiked wild type vs. *tgg*) | Mann-Whitney rank sum test | Amount | *U* = 0.000 | 0.001 |
| | Sequestered allyl glucosinolate per beetle (allyl glucosinolate-spiked wild type vs. *tgg*) | Student's *t*-test | Amount | *t* = 1.689 | 0.119 |
| Long term feeding experiment | Weight, soluble protein and other metabolites (wild type vs. *tgg*) | Student's *t*-test, Mann-Whitney rank sum test | Concentration | see Supplementary Table 5 | |
| | 4MSOB glucosinolate level in beetle gut (wild type vs. *tgg*) | Student's *t*-test | Percentage[A] | *t* = 0.161 | 0.880 |

| Experiment | Comparison | Statistical method | Variable | Statistics | $p$ |
|---|---|---|---|---|---|
| Short-term feeding experiment and inhibition assays | Plant myrosinase inhibition by gut extracts | one-way ANOVA | Activity | $F = 85.639$ | $< 0.01$ |
| Myrosinase activity in feces | Ingested and excreted myrosinase activity | Paired $t$-test[B] | Activity | $t = 10.449$ | $< 0.005$ |
| | Co-incubation of plant myrosinases with feces homogenates | Paired $t$-test | | $t = 0.158$ | 1.000 |
| Supplementary feeding experiment with larvae | 4MSOB glucosinolate in larvae (wild type vs. *tgg*) | Mann-Whitney rank sum test[B] | Percentage[A] | $U = 0.000$ | 0.032 |
| | 4MSOB-cyanide in larvae (wild type vs. *tgg*) | Student's $t$-test[B] | | $t = 17.788$ | 0.004 |
| | 4MSOB isothiocyanate in larvae (wild type vs. *tgg*) | | | $t = 15.341$ | 0.004 |
| | Other 4MSOB isothiocyanate derived metabolites in larvae (wild type vs. *tgg*) | | | $t = 6.300$ | 0.004 |
| | 4MSOB glucosinolate in feces (wild type vs. *tgg*) | | | $t = 2.897$ | 0.080 |
| | 4MSOB-cyanide in larva feces (wild type vs. *tgg*) | | | $t = 2.359$ | 0.184 |
| | 4MSOB isothiocyanate in larva feces (wild type vs. *tgg*) | | | $t = 1.859$ | 0.400 |
| | Other 4MSOB isothiocyanate derived metabolites in larva feces (wild type vs. *tgg*) | | | $t = 1.156$ | 1.000 |
| | 4MSOB glucosinolate and derived metabolites in larvae (wild type vs. *tgg*) | Student's $t$-test, Mann-Whitney rank sum test | Amount | see Supplementary Table 8 | |
| | 4MSOB glucosinolate and derived metabolites in larval feces (wild type vs. *tgg*) | | | see Supplementary Table 8 | |
| | Plant 4MSOB glucosinolate levels (wild type vs. *tgg*) | Student's $t$-test | Concentration | $t = 0.132$ | 0.898 |
| Supplementary long term feeding experiment with larvae | Larval development time (wild type vs. *tgg*) | Mann-Whitney rank sum test | Days | $U = 2{,}319.500$ | 0.420 |
| | Prepupal weight and energy reserves (wild type vs. *tgg*) | Student's $t$-test, Mann-Whitney rank sum test | Fresh weight or concentration | see Supplementary Table 9 | |

Wild type, *Arabidopsis* wild type; *tgg*, *Arabidopsis tgg;* [A]Arcsin-square-root transformed; [B]$p$-values adjusted for false discovery rate in multiple hypothesis testing with Benjamini–Hochberg method

**Supplementary Table 2:** Detected amounts of 4MSOB glucosinolate and derived metabolites in adult *P. armoraciae* beetles and feces after feeding on *Arabidopsis* wild type and *tgg* leave for one day.

| | nmol per individual (mean ± SD; n = 5) | | | | | | | |
|---|---|---|---|---|---|---|---|---|
| | Beetle | | | | Feces | | | |
| Metabolite | wild type-fed | *tgg*-fed | Statistics | *p* | wild type-fed | *tgg*-fed | Statistics | *p* |
| 4MSOB glucosinolate | 3.734 ± 2.862 | 6.229 ± 2.953 | *t* = 1.213 | 0.260 | 0.129 ± 0.166 | 0.867 ± 0.917 | *U* = 5.000 | 0.151 |
| 4MSOB cyanide | 0.483 ± 0.208 | 0.050 ± 0.048 | *U* = 1.000 | 0.016 | 0.809 ± 0.460 | 0.078 ± 0.054 | *U* = 1.000 | 0.016 |
| 4MSOB isothiocyanate | 0.025 ± 0.020 | 0.006 ± 0.002 | *U* = 4.000 | 0.095 | 0.040 ± 0.028 | 0.002 ± 0.001 | *U* = 0.000 | 0.008 |
| Other 4MSOB isothiocyanate-derived metabolites[1] | 0.048 ± 0.025 | 0.008 ±0.004 | *U* = 2.000 | 0.032 | 0.073 ± 0.038 | 0.013 ±0.017 | *t* = 2.882 | 0.020 |
| Total | 4.290 ± 3.058 | 6.292 ± 2.980 | *t* = 0.938 | 0.376 | 1.051 ± 0.639 | 0.960 ± 0.900 | *t* = 0.166 | 0.873 |

[1]comprise 4MSOB isothiocyanate-glutathione conjugate, 4MSOB isothiocyanate-cysteinylglycine conjugate, 4MSOB isothiocyanate-cysteine conjugate. 2-(4-(methylsulfinyl)butylamino)-4,5dihydrothiazole-carboxylic acid, 4MSOB amine, 4MSOB acetamide

**Supplementary Table 3:** Glucosinolate concentration and profile in newly emerged *P. armoraciae* adults reared on *B. rapa.*

| Glucosinolate | nmol × $mg^{-1}$ FW (mean ± SD; n = 20) |
|---|---|
| 3-butenyl | 0.48 ± 0.53 |
| 4-pentenyl | 0.90 ± 0.68 |
| 2-hydroxy-3-butenyl | 3.66 ± 1.65 |
| 2-hydroxy-4-pentenyl | 0.82 ± 0.54 |
| 5-methythiopentyl | 1.53 ± 0.54 |
| benzyl | 0.19 ± 0.14 |
| 2-phenylethyl | 0.35 ± 0.30 |
| indol-3-ylmethyl | 0.23 ± 0.11 |
| 4-methoxyindol-3-ylmethyl | 0.06 ± 0.03 |
| 1-methoxyindol-3-ylmethyl | 0.22 ± 0.08 |
| Total | 8.42 ± 3.26 |

**Supplementary Table 4:** Fresh weight and energy reserves of newly emerged *P. armoraciae* adults after feeding on *Arabidopsis* wild type and *tgg* leaves for 10 days (mean ± SD).

| Parameter | Sex | n | wild type-fed | *tgg*-fed | Statistics | *p* |
|---|---|---|---|---|---|---|
| mg fresh weight | both | 18-19 | 2.56 ± 0.46 | 2.49 ± 0.49 | t = 0.399 | 0.692 |
| | male | 8-9 | 2.12 ± 0.19 | 2.10 ± 0.23 | t = 0.212 | 0.835 |
| | female | 10 | 2.94 ± 0.21 | 2.80 ± 0.42 | t = 0.891 | 0.385 |
| µg soluble protein × mg $FW^{-1}$ | both | 18-19 | 61.35 ± 14.67 | 65.69 ± 15.51 | *U* = 145.000 | 0.438 |
| | male | 8-9 | 52.20 ± 6.65 | 51.36 ± 3.87 | *U* = 34.000 | 0.885 |
| | female | 10 | 69.59 ± 15.03 | 77.15 ± 11.19 | *t* = -1.210 | 0.242 |
| µg total lipids × mg $FW^{-1}$ | both | 18-19 | 3.06 ± 2.70 | 3.86 ± 3.69 | *U* = 151.000 | 0.553 |
| | male | 8-9 | 1.34 ± 1.55 | 1.86 ± 3.16 | *U* = 34.000 | 0.885 |
| | female | 10 | 4.61 ± 2.57 | 5.45 ± 3.29 | *t* = -0.604 | 0.554 |
| µg glycogen × mg $FW^{-1}$ | both | 18-19 | 19.52 ± 5.99 | 21.15 ± 8.82 | *U* = 354.000 | 0.727 |
| | male | 8-9 | 22.07 ± 5.59 | 26.72 ± 10.22 | *U* = 27.000 | 0.413 |
| | female | 10 | 17.22 ± 5.38 | 16.70 ± 3.46 | *t* = 1.099 | 0.286 |
| µg soluble carbohydrates × mg $FW^{-1}$ | both | 18-19 | 3.77 ± 2.13 | 2.93 ± 1.83 | *U* = 132.000 | 0.242 |
| | male | 8-9 | 3.79 ± 2.87 | 2.77 ± 2.12 | *U* = 25.000 | 0.312 |
| | female | 10 | 3.75 ± 1.08 | 3.06 ± 1.53 | *U* = 47.000 | 0.850 |

**Supplementary Table 5:** Protein and metabolite concentrations in rosette leaves of six-week old *Arabidopsis* wild type and *tgg* plants (n = 10; mean ± SD).

| | | wild type | *tgg* | Statistics | *p* |
|---|---|---|---|---|---|
| Soluble protein (mg × g dry weight$^{-1}$) | | 37.010 ±6.028 | 36.805 ± 7.269 | *t* = 0.069 | 0.946 |
| Amino acids (µmol × g dry weight$^{-1}$) | Alanine | 8.040 ± 1.418 | 8.016 ± 1.365 | *t* = 0.038 | 0.970 |
| | Arginine* | 1.498 ± 0.509 | 1.207 ± 0.187 | *U* = 30.000 | 0.140 |
| | Asparagine | 9.540 ± 2.831 | 8.687 ± 1.458 | *U* = 44.000 | 0.678 |
| | Aspartic acid | 6.433 ± 0.764 | 6.862 ± 0.627 | *t* = 1.302 | 0.209 |
| | Glutamic acid | 49.035 ± 5.004 | 50.804 ± 4.426 | *t* = 0.794 | 0.437 |
| | Glutamine | 100.721 ± 17.043 | 109.767 ± 13.346 | *t* = 1.254 | 0.226 |
| | Histidine* | 2.741 ± 0.341 | 2.909 ± 0.392 | *t* = 0.968 | 0.346 |
| | Isoleucine* | 1.100 ± 0.188 | 1.041 ± 0.093 | *t* = 0.847 | 0.408 |
| | Leucine* | 0.959 ± 0.128 | 0.898 ± 0.134 | *t* = 0.991 | 0.335 |
| | Lysine* | 0.781 ± 0.088 | 0.747 ± 0.082 | *t* = 0.836 | 0.414 |
| | Methionine* | 0.253 ± 0.022 | 0.237 ± 0.035 | *t* = 1.185 | 0.252 |
| | Phenylalanine* | 0.800 ± 0.226 | 0.727 ± 0.092 | *U* = 43.000 | 0.623 |
| | Proline | 3.984 ± 1.007 | 4.906 ± 1.017 | *U* = 18.000 | 0.017 |
| | Serine | 16.839 ± 2.716 | 16.025 ± 0.996 | *U* = 45.000 | 0.734 |
| | Threonine* | 11.012 ± 1.184 | 11.675 ± 0.850 | *t* = 1.365 | 0.189 |
| | Tryptophane* | 0.139 ± 0.039 | 0.130 ± 0.016 | *U* = 46.000 | 0.791 |
| | Tyrosine | 0.457 ± 0.087 | 0.392 ± 0.042 | *t* = 2.013 | 0.059 |
| | Valine* | 1.820 ± 0.265 | 1.764 ± 0.128 | *t* = 0.562 | 0.581 |
| | Total essential amino acids* | 21.103 ± 2.171 | 21.336 ± 1.176 | *t* = 0.283 | 0.781 |
| | Total non-essential amino acids | 185.508 ± 23.637 | 196.770 ± 15.005 | *t* = 1.207 | 0.243 |
| | Total amino acids | 206.611 ± 24.816 | 218.107 ± 15.327 | *t* = 1.182 | 0.252 |
| Soluble sugars (mg × g dry weight$^{-1}$) | Glucose | 0.154 ± 0.045 | 0.152 ± 0.044 | *t* = 0.090 | 0.929 |
| | Fructose | 0.069 ± 0.019 | 0.049 ± 0.011 | *t* = 2.796 | 0.012 |
| | Sucrose | 0.125 ±0.027 | 0.145 ± 0.016 | *t* = 2.186 | 0.042 |
| | Raffinose | 0.080 ± 0.021 | 0.087 ± 0.009 | *U* = 45.000 | 0.307 |
| | Total sugars | 0.428 ± 0.079 | 0.433 ± 0.060 | *t* = 0.538 | 0.597 |
| Glucosinolates (µmol × g dry weight$^{-1}$) | 3MSOP glucosinolate | 1.924 ± 0.251 | 1.834 ± 0.130 | *U* = 38.000 | 0.385 |
| | 4MSOB glucosinolate | 15.071 ± 1.537 | 14.811 ± 1.253 | *t* = 0.393 | 0.699 |
| | 5MSOP glucosinolate | 0.510 ± 0.047 | 0.513 ± 0.042 | *t* = 0.146 | 0.885 |
| | 7MSOH glucosinolate | 0.249 ± 0.032 | 0.231 ± 0.016 | *t* = 1.543 | 0.140 |
| | 8MSOO glucosinolate | 1.209 ± 0.094 | 1.263 ± 0.082 | *t* = 1.298 | 0.211 |
| | 4MTB glucosinolate | 1.200 ± 0.132 | 1.183 ± 0.126 | *t* = 0.285 | 0.779 |
| | 4OHI3M glucosinolate | 0.022 ± 0.011 | 0.016 ± 0.010 | *t* = 1.216 | 0.240 |
| | I3M glucosinolate | 1.756 ± 0.218 | 1.532 ± 0.138 | *t* = 2.598 | 0.018 |
| | 4MOI3M glucosinolate | 0.394 ± 0.035 | 0.368 ± 0.039 | *U* = 28.000 | 0.104 |
| | 1MOI3M glucosinolate | 0.088 ± 0.035 | 0.127 ± 0.105 | *U* = 37.000 | 0.345 |
| | Total glucosinolates | 22.422 ± 1.902 | 21.877 ± 1.414 | *t* = 0.689 | 0.500 |

*, essential amino acids; 3MSOP, 3-methylsulfinylpropyl; 4MSOB, 4-methylsulfinylbutyl; 5MSOP, 5-methylsulfinylpentyl; 7MSOH, 7-methylsulfinylheptyl; 8MSOO, 8-methylsulfinyloctyl; 4MTB, 4-methylthiobutyl; 4OHI3M, 4-hydroxyindol-3-ylmethyl; I3M, indol-3-ylmethyl; 4MOI3M, 4-methoxyindol-3-ylmethyl; 1MOI3M, 1-methoxyindol-3-ylmethyl;

**Supplementary Table 6:** Distribution of 4MSOB glucosinolate in the gut and remaining body of *P. armoraciae* adults after 1 min feeding and 5 min digestion.

| Treatment | glucosinolate proportion (mean % ± SE; n = 3) | | | |
|---|---|---|---|---|
| | Gut | Rest of body | *t*[1] | *p* |
| wild type-fed | 16.5 ± 2.6 | 83.5 ± 2.6 | 8.529 | 0.013 |
| *tgg*-fed | 16.1 ± 2.2 | 83.9 ± 2.2 | 10.103 | 0.010 |

[1]Data were arcsine square root transformed before statistical analysis by Student's *t*-test

**Supplementary Table 7:** *Arabidopsis* TGG1-derived peptides detected by nano-UPLC-$MS^E$ in feces of *P. armoraciae* adults fed with on *Arabidopsis* wild type leaves.

| Amino acid sequence | sample number[1] | Sequence identity |
|---|---|---|
| LFNSGNFEK | 20 | TGG1 |
| GFIFGVASSAYQVEGGR | 20 | TGG1 |
| GLNVWDSFTHR | 20 | TGG1 |
| GGADLGNGDTTCDSYTLWQK | 19 | TGG1 |
| FSIAWSR | 19, 20 | TGG1 and TGG6 |
| YYNGLIDGLVAK | 19, 20 | TGG1 |
| NWITINQLYTVPTR | 19, 20 | TGG1 |
| GYALGTDAPGR | 19, 20 | TGG1 and TGG2 |
| DDQKGMIGPVMITR* | 19 | TGG1 |
| GMIGPVMITR | 19, 20 | TGG1 |
| WFLPFDHSQESK | 19, 20 | TGG1 |
| LPEFSETEAALVK | 19, 20 | TGG1 |
| GIYYVMDYFK | 19, 20 | TGG1 |
| TTYGDPLIYVTENGFSTPGDEDFEK | 20 | TGG1 |

[1]corresponding to sample number in Figure 4A; *contains one missed tryptic cleavage site

**Supplementary Table 8:** Detected amounts of 4MSOB glucosinolate and derived metabolites in *P. armoraciae* larvae and feces after feeding on *Arabidopsis* wild type and *tgg* leaves after one day feeding.

| | nmol per individual (mean ± SD; n = 5) | | | | | | | |
|---|---|---|---|---|---|---|---|---|
| | Larva | | | | Feces | | | |
| Metabolite | wild type-fed | *tgg*-fed | Statistics | *p* | wild type-fed | *tgg*-fed | Statistics | *p* |
| 4MSOB glucosinolate | 0.001 ± 0.001 | 0.372 ± 0.290 | $t$ = 2.558 | 0.034 | 0.100 ± 0.078 | 0.215 ± 0.165 | $t$ = 1.262 | 0.242 |
| 4MSOB cyanide | 0.684 ± 0.415 | 0.207 ± 0.130 | $U$ = 5.000 | 0.151 | 1.158 ± 0.628 | 0.537 ± 0.342 | $t$ = 1.737 | 0.121 |
| 4MSOB isothiocyanate | 0.016 ± 0.008 | 0.728 ± 0.286 | $U$ = 0.000 | 0.008 | 0.013 ± 0.007 | 0.004 ± 0.003 | $t$ = 2.231 | 0.056 |
| Other 4MSOB isothiocyanate-derived metabolites[1] | 0.023 ± 0.011 | 0.334 ± 0.162 | $U$ = 0.000 | 0.008 | 0.051 ± 0.027 | 0.027 ± 0.030 | $t$ = 1.193 | 0.267 |
| Total | 0.724 ± 0.430 | 1.641 ± 0.763 | $t$ = 2.093 | 0.070 | 1.322 ± 0.709 | 0.784 ± 0.506 | $t$ = 1.236 | 0.251 |

[1]comprise 4MSOB isothiocyanate-glutathione conjugate, 4MSOB isothiocyanate-cysteinylglycine conjugate, 4MSOB isothiocyanate-cysteine conjugate, 2-(4-(methylsulfinyl)butylamino)-4,5dihydrothiazole-carboxylic acid, 4MSOB amine, 4MSOB acetamide

**Supplementary Table 9:** Fresh weight and energy reserves of *P. armoraciae* prepupae reared on *A. thaliana* wild type and *tgg* leaves from the early second instar. Mean ± SD are listed.

| Parameter | n | wild type-fed | *tgg*-fed | Statistics | *p* |
|---|---|---|---|---|---|
| developmental time (days) | 68-74 | 33 ± 2 | 34 ± 3 | $U$ = 2,319.500 | 0.420 |
| mg FW × prepupa$^{-1}$ | 68-74 | 1.34 ± 0.23 | 1.32 ± 0.22 | $t$ = 0.331 | 0.741 |
| µg soluble protein × mg FW$^{-1}$ | 11 | 72.78 ± 6.91 | 77.47 ± 13.06 | $U$ = 49.000 | 0.470 |
| µg total lipids × mg FW$^{-1}$ | 11 | 51.87 ± 3.42 | 41.77 ± 3.46 | $t$ = -1.586 | 0.129 |
| µg glycogen × mg FW$^{-1}$ | 11 | 157.58 ± 34.98 | 128.68 ± 17.56 | $U$ = 52.000 | 0.599 |
| µg soluble carbohydrates × mg FW$^{-1}$ | 11 | 55.52 ± 2.88 | 51.24 ± 4.96 | $t$ = 0.526 | 0.605 |

**SUPPLEMENTARY FIGURES**

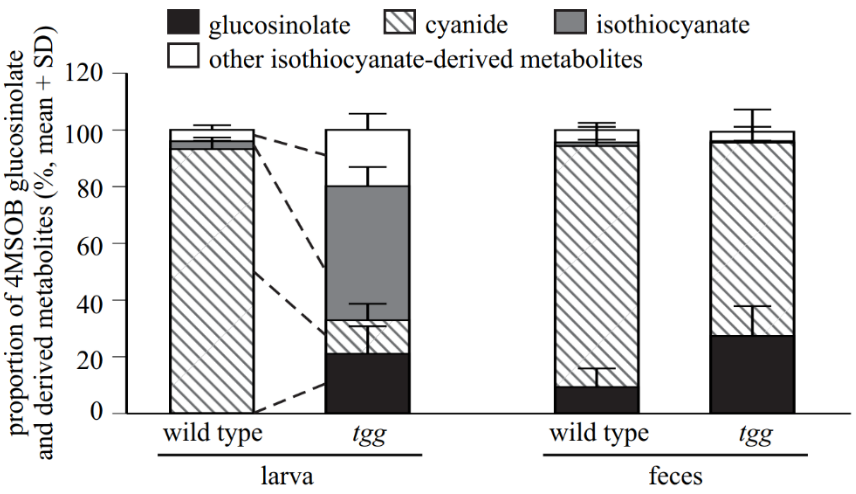


**Supplementary Figure 1:** Proportions of 4-methylsulfinylbutyl (4MSOB) glucosinolate and derived hydrolysis products detected in the body and feces of *P. armoraciae* larvae fed with *Arabidopsis* wild type or myrosinase-deficient *tgg* leaves (n = 5). Glucosinolates and hydrolysis products were extracted with 50% methanol and analyzed by LC-MS/MS. Detected amounts of metabolites were expressed relative to the total amounts of all detected metabolites in larvae or feces (set to 100%). Dashed lines indicate significant differences between samples ($p < 0.05$). Statistical results are shown in Supplementary Table 1. 4MSOB cyanide corresponds to the nitrile formed from 4MSOB glucosinolate. Other isothiocyanate-derived metabolites comprise 4MSOB isothiocyanate-glutathione conjugate, 4MSOB isothiocyanate-cysteinylglycine conjugate, 4MSOB isothiocyanate-cysteine conjugate, 2-(4-(methylsulfinyl)butylamino)-4,5dihydrothiazole-carboxylic acid, 4MSOB amine, and 4MSOB acetamide.

154
